# Exploring CDK4/6-Dependencies in *ex vivo* Ovarian Cancer Models

**DOI:** 10.1101/2025.07.11.664318

**Authors:** Jake McDonald-Pike, Camilla Coulson-Gilmer, Samantha Littler, Bethany M. Barnes, James Altringham, Anthony Tighe, Joanne C. McGrail, Stephen S. Taylor

## Abstract

Ovarian cancer (OC) is a clinically and molecularly heterogeneous disease with limited treatment options for the majority of patients, particularly those with homologous-recombination-proficient high-grade serous ovarian cancer (HGSOC) and rarer subtypes such as low-grade serous ovarian cancer. Deregulation of the G1/S cell cycle network is common across all subtypes, suggesting subtype-agnostic vulnerabilities. Here, we assessed CDK4/6 dependency using the selective inhibitor palbociclib across 20 patient-derived *ex vivo* OC models. A subset of models, including four HGSOC and six rarer subtypes, exhibited marked sensitivity to palbociclib, characterised by low *CDKN2A*/*CDKN2B* expression, Rb hypophosphorylation, and G1 cell cycle arrest. In contrast, resistant models showed high *CDKN2A* expression and reduced or absent *RB1*. Notably, *ABCB1* overexpression—a known resistance mechanism in OC—did not mediate palbociclib resistance. Analysis of longitudinal models revealed diminished CDK4/6 dependency following treatment, accompanied by increased *CDKN2A* expression. These findings support a model of G1/S control in which tumours diverge into CDK4/6- or CDK2-driven proliferation states, with *CDKN2A* as a potential biomarker to guide patient selection. The predominance of CDK4/6-inhibitor-resistant HGSOC highlights a priority population for CDK2-targeted therapies, offering new treatment strategies for patients with otherwise limited options.

## Introduction

Ovarian cancer (OC) is the deadliest gynaecological malignancy and the sixth leading cause of cancer-related death among women, with 10-year survival rates of approximately 35% (Cancer Research UK., 2025). OC is a clinically, histologically, and molecularly heterogeneous disease comprising multiple distinct subtypes (Prat, 2012; Hollis, 2023). High-grade serous ovarian cancer (HGSOC) is the most prevalent and lethal subtype, typically diagnosed at an advanced stage (Lisio et al., 2019). While most HGSOC patients initially respond well to treatment—usually neoadjuvant carboplatin-paclitaxel chemotherapy followed by surgery and adjuvant chemotherapy—relapse with drug-resistant disease is common (Bowtell et al., 2015). Despite recent advances, including anti-angiogenic agents and PARP1/2 inhibitors for homologous recombination (HR)-deficient disease, treatment options for the majority of HGSOC patients remain limited (González-Martín et al., 2023; Morgan et al., 2023).

Rarer subtypes—including endometrioid (ENOC), low-grade serous (LGSOC), clear cell (CCOC), and mucinous ovarian cancer (MOC)—also contribute significantly to overall disease burden. These subtypes tend to exhibit intrinsic chemotherapy resistance and remain underserved by subtype-specific therapies (Hollis, 2023). Together, HR-proficient HGSOC and rarer OC subtypes represent an important unmet clinical need, underscoring the urgency of exploring targeted therapeutic strategies.

One emerging area for therapeutic exploration is the G1/S cell cycle control network, which is frequently deregulated across OC subtypes, leading to uncoupling of proliferation from mitogenic signalling. In normal cells, G1–S progression is tightly regulated by cyclin-dependent kinases (CDKs) (Hume et al., 2020; Rubin et al., 2020). Mitogenic cues induce D-type cyclins, which activate CDK4/6, leading to phosphorylation of retinoblastoma protein (Rb), release of E2F transcription factors, induction of cyclin E and activation of CDK2. This cascade drives S-phase entry and is restrained by tumour suppressors such as p53, and CDK inhibitors including p16 (*CDKN2A*) and p21 (*CDKN1A*).

In HGSOC, near-universal *TP53* mutation disables G1 checkpoint control, permitting unchecked proliferation despite widespread genomic instability (Ahmed et al., 2010). The Rb pathway is disrupted in approximately two-thirds of cases (TCGA, 2011), most commonly via *CDKN2A* downregulation or deletion (∼32%), *CCNE1/2* amplification (∼20%), or *RB1* loss (∼10%). These alterations promote aberrant E2F activity and are often reinforced by amplification of *CDK2/4/6*, *E2F1/2/3*, or *MYC*, collectively driving uncontrolled S-phase entry. Parallel activation of mitogenic pathways occurs in ∼45% of HGSOC, frequently due to loss of upstream suppressors such as *NF1* or *PTEN*, or amplification of downstream effectors including *PIK3CA*, *KRAS*, and *AKT1/2*, leading to cyclin D upregulation (∼15%). Similar signalling alterations are common in rarer subtypes, with frequent *KRAS*, *BRAF*, and *PIK3CA* mutations (Singer et al., 2003; Hunter et al., 2015; Friedlander et al., 2016; Cheasley et al., 2019; Pierson et al., 2020; Cheasley et al., 2021; Iida et al., 2021). *CDKN2A* is also frequently deleted in LGSOC (Hunter et al., 2015; Cheasley et al., 2021) and altered in ∼50% of MOC cases (Cheasley et al., 2019). Despite subtype-specific features, these alterations converge on persistent mitogenic signalling and G1/S checkpoint disruption.

Crucially, these events generate distinct dependencies on specific cyclin–CDK complexes. Functional studies have shown that tumours with intact Rb and *CDKN2A* loss or *CCND1* amplification are often dependent on CDK4/6 for G1 progression (Knudsen et al., 2022; Zhang et al., 2022). In contrast, tumours with *CCNE1* amplification or *RB1* loss bypass CDK4/6 and rely instead on CDK2 (Sine et al., 2025). This functional divergence suggests that G1/S deregulation constitutes a context-specific vulnerability that may be therapeutically exploited using selective CDK inhibitors.

Selective CDK4/6 inhibitors (palbociclib, ribociclib, abemaciclib) are now standard-of-care for ER-positive, HER2-negative advanced breast cancer, and their clinical success has motivated exploration in other malignancies (Wang et al., 2024). Given that Rb remains intact in most OC cases and that many harbour alterations suggestive of CDK4/6 dependency, CDK4/6 inhibition represents a promising strategy for a defined subset of OC patients. Indeed, preclinical studies have shown that CDK4/6 inhibitors can be effective in both HGSOC and rarer subtypes, with sensitivity often associated with Rb proficiency and low *CDKN2A* expression (Konecny et al., 2011; Taylor-Harding et al., 2015; Dall’Acqua et al., 2017; Iyengar et al., 2018; Yi et al., 2019). However, these studies have largely relied on established cell lines, which do not fully recapitulate the molecular diversity or complexity of primary tumours (Domcke et al., 2013; Barnes et al., 2021). There is therefore a pressing need to assess CDK4/6 inhibition in more clinically relevant systems.

To address this, we leveraged a Living Biobank of *ex vivo* patient-derived ovarian cancer models (OCMs) that retain the molecular and phenotypic features of primary tumours (Nelson et al., 2020; Nelson et al., 2023). This resource includes chemo-naïve, post-treatment, and longitudinal models from individual patients. Their robust proliferative capacity and experimental tractability make them well suited for functional drug screening. In this study, we assess the efficacy of CDK4/6 inhibition using palbociclib across a diverse panel of 20 *ex vivo* OCMs. We quantify drug sensitivity using long-term colony formation assays and explore how molecular alterations in the G1/S network inform response to CDK4/6 inhibition.

## Results

### Establishing an assay to determine palbociclib sensitivity

To explore CDK4/6 inhibitor responses in a Living Biobank of *ex vivo* ovarian cancer models (OCMs), we first established a drug screening assay designed to (a) capture biologically meaningful drug-induced anti-proliferative effects and (b) confirm the on-target, cytostatic activity of palbociclib. To optimise assay conditions, we selected RKO cells—a human colorectal cancer line previously shown in-house to exhibit reduced proliferation upon palbociclib treatment (data not shown). To measure anti-proliferative effects, we initially deployed colony formation assays (CFAs). RKO cells were treated with increasing concentrations of palbociclib, incubated for up to three weeks with weekly media and drug replenishment, then fixed, stained, and imaged (Figure 1a). Bound stain was extracted and quantified to generate dose–response curves. As expected, palbociclib inhibited colony formation in a concentration-dependent manner, with an estimated GI₅₀ of ∼150 nM (Figure 1b–c), consistent with reported sensitivities in other palbociclib-responsive cell lines (Konecny et al., 2011).

**Figure 1.**
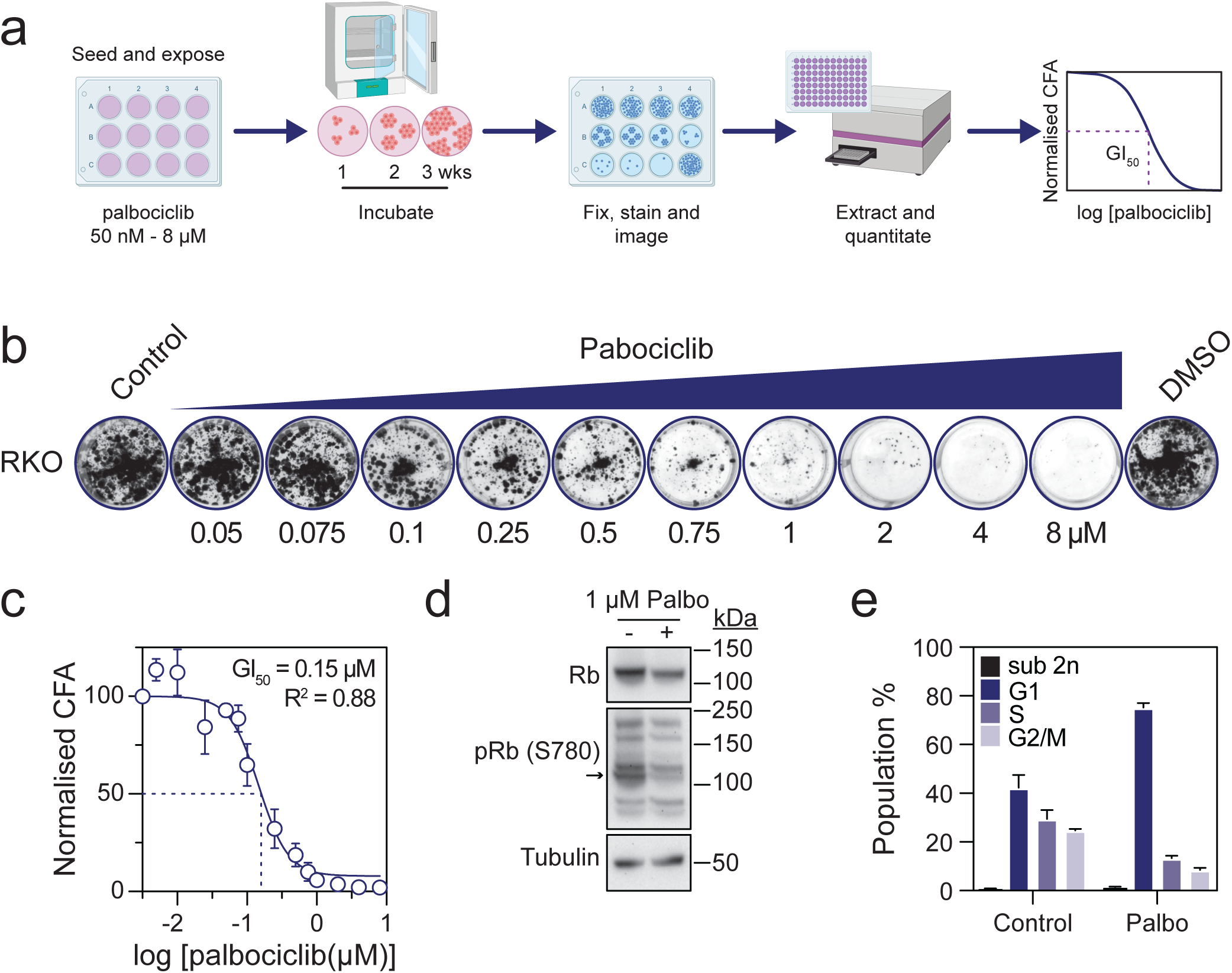
Assay to determine palbociclib sensitivity a. Schematic overview of colony formation assay (CFA) to determine palbociclib GI_50_ values for cell lines and ovarian cancer models (OCMs). b. CFA of RKO cells exposed to increasing concentrations of palbociclib. c. XY graph quantitating CFA values across a range of palbociclib concentrations. Each value represents the mean ± s.e.m. from at least three independent experiments. The dose response curve was calculated using a non-linear regression (log(inhibitor) vs. response – variable slope (four parameters)) to determine the GI_50_ value and goodness of fit (R^2^). d. Immunoblot of RKO cells treated with 1 µM palbociclib showing reduced Rb phosphorylation (pRb, serine 780). pRb is marked with an arrow. Total Rb and tubulin were used as loading controls. Blot shown is a representative biological replicate. e. Bar graph of flow cytometry data from RKO cells exposed to 1 µM palbociclib for 24 hours. DNA content histograms were modelled to determine the percentage of cells in G1, S-phase (S) and G2 or mitosis (G2/M). Cells with sub-2n DNA contents were also quantified.

To determine whether the observed anti-proliferative effects were consistent with on-target cytostatic activity of palbociclib, we assessed Rb phosphorylation and cell cycle distribution in RKO cells treated with 1 μM palbociclib for 24 hours. Immunoblotting confirmed that Rb was expressed in both untreated and treated cells (Figure 1d), with a modest reduction in total Rb levels following treatment, possibly reflecting decreased protein stability. Crucially, phosphorylation of Rb at serine 780—a known CDK4/6 target site—was markedly reduced in palbociclib-treated cells, consistent with effective CDK4/6 inhibition. Flow cytometric analysis was also consistent with palbociclib-induced G1 arrest, with the G1 population increasing from 42.3% to 75.2%, accompanied by corresponding reductions in S and G2/M phase cells (Figure 1e). The sub-G1 population remained low in both conditions, indicating minimal apoptosis. Together, these findings are consistent with palbociclib exerting an on-target, cytostatic effect in RKO cells, inhibiting Rb phosphorylation and inducing G1 cell cycle block. Collectively, these data validate the CFA as a robust assay to capture biologically relevant, anti-proliferative responses to CDK4/6 inhibition.

### A subset of patient-derived ovarian cancer models are sensitive to palbociclib

Having established a CFA to measure CDK4/6 inhibitor responses, we next evaluated the effects of palbociclib across a panel of patient-derived *ex vivo* ovarian cancer models (OCMs) (Nelson et al., 2020; Nelson et al., 2023). For this pilot screen, we selected 15 OCMs encompassing HGSOC, LGSOC, and CCOC subtypes (Table S1). Upon palbociclib treatment, we observed a spectrum of responses, ranging from pronounced sensitivity to marked resistance (Figure S1). For example, OCM.314 was markedly inhibited at 100 nM and was completely ablated at 8 μM, whereas OCM.149 remained largely unaffected even at 4 μM. Dose–response curves and GI₅₀ values confirmed this variation in sensitivity (Figure 2a). Six OCMs showed concentration-dependent reductions in colony formation at sub-micromolar concentrations, consistent with sensitivity to palbociclib. Notably, OCM.314 (CCOC) and OCM.292 (HGSOC) were particularly sensitive, with GI₅₀ values of 60 nM and 100 nM, respectively. In contrast, nine OCMs showed minimal response at concentrations up to 4 μM, indicating resistance. In several cases, inhibition of colony formation was only evident at concentrations exceeding 4 μM—a range where palbociclib is known to exert off-target effects (Fry et al., 2004). Across the panel, GI₅₀ values <1 μM were indicative of sensitivity, whereas values ≥6 μM reflected resistance. Applying these thresholds, we classified six of the 15 OCMs as sensitive and nine as resistant to palbociclib.

**Figure 2.**
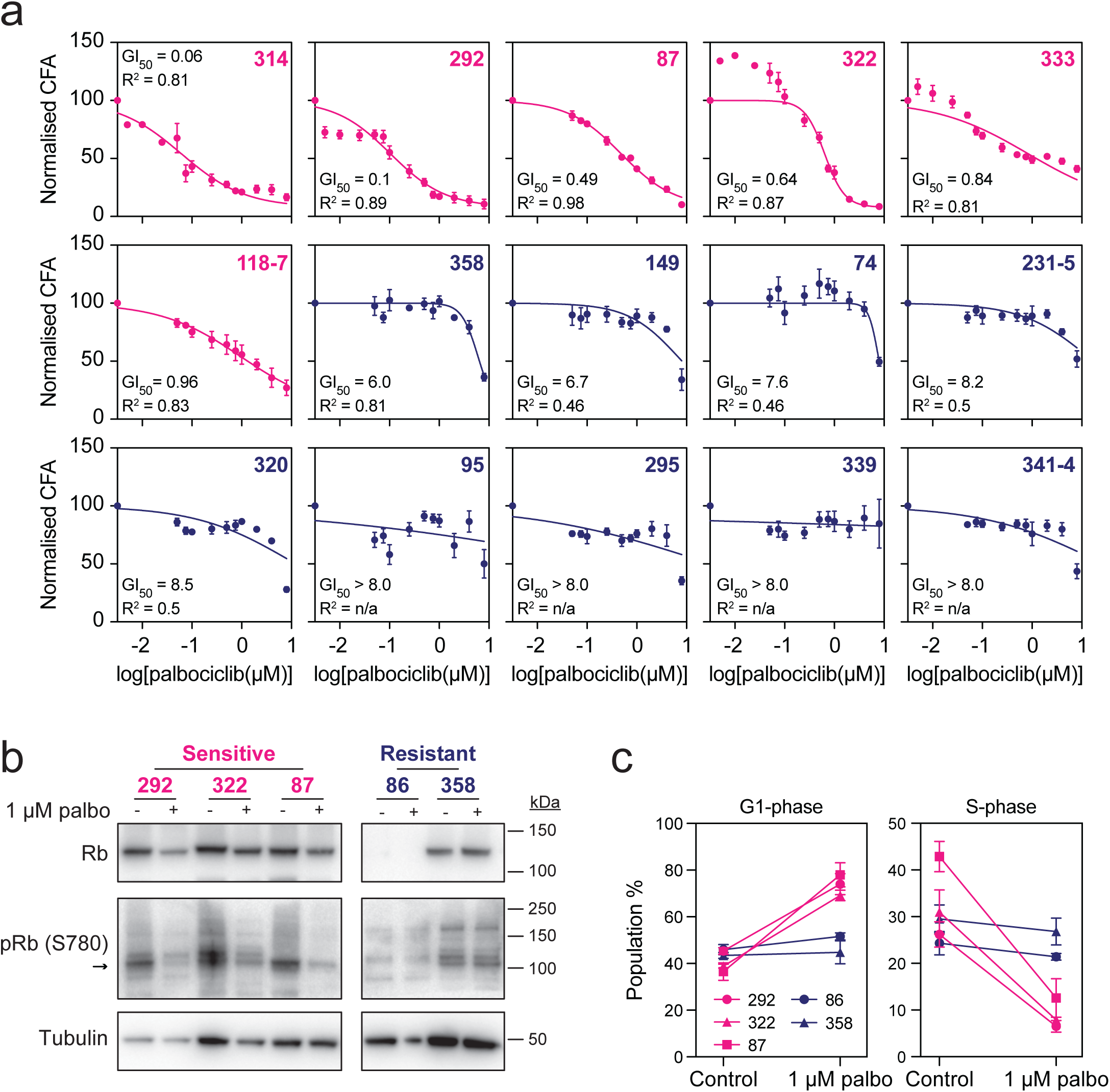
Palbociclib inhibits a subset of patient-derived ovarian cancer models a. XY graph quantitating CFA values across a panel of 15 patient-derived ovarian cancer models (OCMs) in response to a range of palbociclib concentrations. Each value represents the mean ± s.e.m. from at least three independent experiments. Dose response curves were generated to determine GI_50_ values, with R^2^ values indicating the goodness of fit for the non-linear regression. The OCMs are arranged top-left to bottom-right in order of increasing GI_50_. Six OCMs with GI_50_ values < 1µM were classified as sensitive (magenta) while nine OCMS with GI_50_ values ≥6 µM were classified as resistant (dark blue). b. Immunoblot of five OCMs cells treated with 1 µM palbociclib, processed to detect total Rb and phospho-Rb (serine 780, arrow). Tubulin was used as a loading control. Blot shown is representative of multiple biological replicates. Palbociclib supresses pRb in three sensitive OCMs (292, 322 and 87, magenta). In the two resistant OCMs (87 and 358, dark blue), Rb is not detected in OCM.86 and no decrease in pRb is detected for OCM.358. c. Before-and-after graphs quantitating the percentage of cells in G1 and S-phase for five OCMs in response to 1 µM palbociclib for 24 hours. Each value shows the mean ± s.e.m. from two independent experiments. The percentage of G1 and S-phase cells increases and decreases respectively in the sensitive OCMs (magenta) relative to the resistant (dark blue).

To confirm that the observed responses to palbociclib in OCMs were consistent with CDK4/6 inhibition, we examined Rb phosphorylation (Figure 2b) and cell cycle progression (Figure 2c). In three palbociclib-sensitive models—OCMs 292, 322, and 87—treatment led to a marked reduction in phospho-Rb levels. As in RKO cells (Figure 1d), we also observed a modest reduction in total Rb. In contrast, two resistant models showed no change in phospho-Rb following palbociclib exposure. Although immunoreactive bands were detected, we suspect these represent non-specific cross-reactive species. Indeed, in OCM.86, Rb was undetectable by pan-Rb antibody (Figure 2b), and RNA-seq analysis confirmed the absence of *RB1* expression (see below), indicating this model is derived from an Rb-null tumour. In OCM.358, total Rb was present but phospho-Rb signal remained unchanged following treatment. These findings suggest that the reduction in Rb phosphorylation observed in sensitive models is a consequence of CDK4/6 inhibition, while the modest decrease in total Rb likely reflects suppressed proliferation.

Consistent with these findings, DNA content analysis revealed G1-phase accumulation and a corresponding reduction in S-phase cells in palbociclib-sensitive models (Figure 2c). In contrast, palbociclib-resistant models showed minimal changes in cell cycle distribution. Collectively, these results indicate that the anti-proliferative effects observed in palbociclib-sensitive OCMs are mediated by on-target inhibition of CDK4/6, resulting in suppressed Rb phosphorylation and G1 cell cycle arrest.

### Palbociclib resistance is not mediated by MDR1-dependent drug eflux

To explore mechanisms underlying differential palbociclib sensitivity, we first investigated drug-efflux transporters. *ABCB1*, which encodes MDR1/P-glycoprotein—the archetypal ATP-dependent efflux pump—is a well-established driver of clinical drug resistance in HGSOC. It is frequently overexpressed in recurrence, often via promoter-swapping translocations in exon 1 (Patch et al., 2015). One study found that ∼18% of recurrent HGSOC cases overexpressed *ABCB1* (Christie et al., 2019). Consistent with this, we recently demonstrated that a substantial subset of OCMs overexpress *ABCB1*, leading to paclitaxel resistance that can be reversed by co-treatment with elacridar, a selective MDR1 inhibitor (Tighe et al., 2025). Notably, these OCMs were derived from patients who had received high cumulative doses of paclitaxel in the clinic. Efflux-mediated resistance can also confound screens with novel therapeutics. We have shown that overexpression of *ABCB1* and *ABCG2* confers resistance to the SUMO-activating enzyme inhibitor ML-792, and that co-exposure to elacridar unmasks SUMO-signalling dependencies in additional OCMs (Littler et al., 2025). Thus accounting for ABC transporter–mediated drug efflux is important when interpreting drug sensitivity in our Living Biobank.

Interrogation of RNA sequencing (RNA-seq) data associated with the biobank (Nelson et al., 2020; Barnes et al., 2021; Coulson-Gilmer et al., 2021; Littler et al., 2025; Tighe et al., 2025), revealed that palbociclib-resistant HGSOC OCMs overexpressed *ABCB1* (Figure S2a). And indeed, OCMs 74-1, 95, 149, 339, and 358 are taxol-sensitised when co-exposed to elacridar (Tighe et al., 2025). To test whether this overexpression contributed to palbociclib resistance, we treated four high-*ABCB1* models (OCMs 74-1, 95, 149, and 358) with palbociclib in the presence or absence of elacridar. However, co-exposure to elacridar did not enhance sensitivity to palbociclib (Figure S2b), suggesting that palbociclib is not a substrate for MDR1.

To confirm this directly, we used an RKO derivative carrying a tetracycline-inducible MDR1 transgene and a GFP-tagged histone for live-cell proliferation tracking (Tighe et al., 2025). Tetracycline induction of MDR1 reversed the anti-proliferative effects of paclitaxel but had no impact on palbociclib sensitivity, as assessed by fluorescence time-lapse microscopy and colony formation assays across a range of palbociclib doses (Figure S2c, d). In summary, these data demonstrate that palbociclib is not a good MDR1 substrate and that *ABCB1* overexpression does not underlie palbociclib resistance in our OCM panel.

### Identification of additional palbociclib-resistant models based on *CKDN2A* expression

To explore alternative drivers of differential palbociclib responses, we turned our attention to the G1/S cell cycle control network (Figure 3a), and interrogated RNA-seq data to explore the expression of 20 G1/S genes. Examining relative expression levels for the 15 OCMs analysed in the pilot screen (Figure 3b, left panel), did not reveal any overt clustering patterns. However, we did observe that the nine resistant models exhibited relatively high expression of *CDKN2A* and *CDKN2B,* which encode the INK4 family inhibitors p16/INK4A and p15/INK4B. Note that the *CDKN2A* locus also encodes p14/ARF. Conversely, the six sensitive OCMs showed marginally higher *RB1* expression.

**Figure 3.**
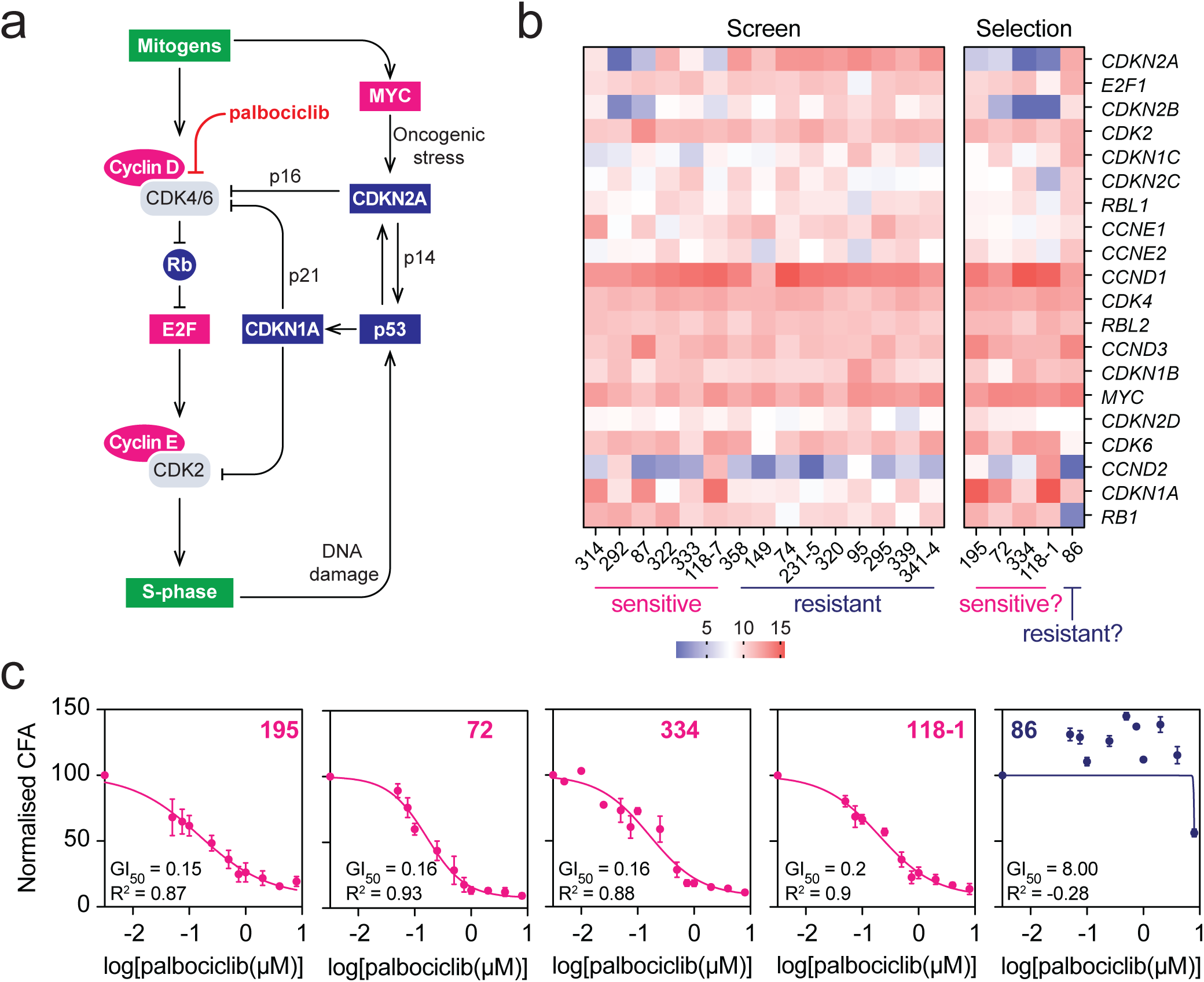
Identification of additional palbociclib-resistant models based on *CKDN2A* expression a. Network diagram describing G1/S cell cycle controls whereby mitogens activate Cyclin-D-CDK4/6 complexes. This inhibits Rb, alleviating inhibition of E2F transcription factors resulting in activation of Cyclin-E-CDK2 complexes which drive the cell towards S-phase. In response to DNA damage or oncogenic stress (e.g. via persistent MYC activation), p53 activation induces *CDKN1A* and *CDKN2A,* which encode Cyclin-dependent Kinase Inhibitors (CKIs) from the CIP/KIP and INK4 families, respectively. While p21 can inhibit CDK4/6 and CDK2, p16 only inhibits CDK4/6. Palbociclib directly blocks CDK4/6 activity. b. Heatmap showing the relative expression levels of 20 G1/S regulator genes across 20 OCMs, with red and blue indicating relatively high and low expression, respectively. The left-hand block shows the 15 OCMs screened in Figure 1, partitioned into six sensitive (left) and nine resistant (right) OCMs. The right-hand block shows five additional OCMs selected on the basis of *CDKN2A* expression, four with relatively low levels and one with relatively high levels. c. XY graph quantitating CFA values across the panel of five selected OCMs, in response to a range of palbociclib concentrations. Each value represents the mean ± s.e.m. from at least three independent experiments. Dose response curves were generated to determine GI_50_ values, with the R^2^ values indicating the goodness of fit. The OCMs are arranged left to right in order of increasing GI_50_. Four OCMs with low *CDKN2A* levels are sensitive, with GI_50_ values < 1µM (195, 72, 334 and 118-1, magenta). OCM.86, which has high *CDKN2A*, is resistant, with a GI_50_ value ≥6 µM (dark blue).

Given reports linking Rb proficiency and low *CDKN2A* expression with CDK4/6-dependency (Knudsen et al., 2022; Zhang et al., 2022), we hypothesised that OCMs with low INK4 levels and intact Rb would be palbociclib-sensitive. We therefore explored the RNA-seq data for OCMs that matched this molecular profile, identifying four additional candidates—OCMs 195, 72, 334, and 118-1—all of which exhibited low *CDKN2A*/*CDKN2B* and high *RB1* expression (Figure 3b, right panel). We also selected OCM.86, which displayed high *CDKN2A* and low *RB1*, as a predicted palbociclib-resistant model. Using our CFA, we tested these five OCMs. As predicted, OCMs 195, 72, 334, and 118-1 were sensitive to low micromolar palbociclib concentrations, each with GI₅₀ < 1 µM (Figure 3c and Figure S3). In contrast, OCM.86 showed only a modest reduction in colony formation at 8 µM, failing to yield a meaningful GI₅₀. These results confirm that palbociclib sensitivity in OCMs correlates with low *CDKN2A*/*CDKN2B* and high *RB1*, and suggest that this expression signature may be sufficient to predict CDK4/6 inhibitor responsiveness in ovarian cancer.

### *CDKN2A* is overexpressed in palbociclib-resistant ovarian cancer models

Armed with a panel of 20 OCMs—10 palbociclib-sensitive and 10 resistant (Figure 4a)—we next compared their clinical and molecular characteristics. All 10 resistant models were derived from HGSOC (Figure 4b; Table S1). In contrast, the 10 sensitive OCMs comprised three LGSOC, two CCOC, one MOC, and four additional HGSOC models. Consistent with our thresholds, sensitive models exhibited GI₅₀ values less than1 µM, whereas resistant models displayed GI₅₀ values greater than 6 µM. For four of the most resistant models, dose–response curves did not yield a definable GI₅₀, so we assigned them a nominal value of 8 µM (Figure 4b). Moreover, six of the sensitive OCMs were highly responsive, with GI₅₀ values less than 200 nM. This clear partitioning by GI₅₀ threshold was statistically significant (Figure 4c), reinforcing the distinct sensitivity versus resistance phenotypes in our cohort.

**Figure 4.**
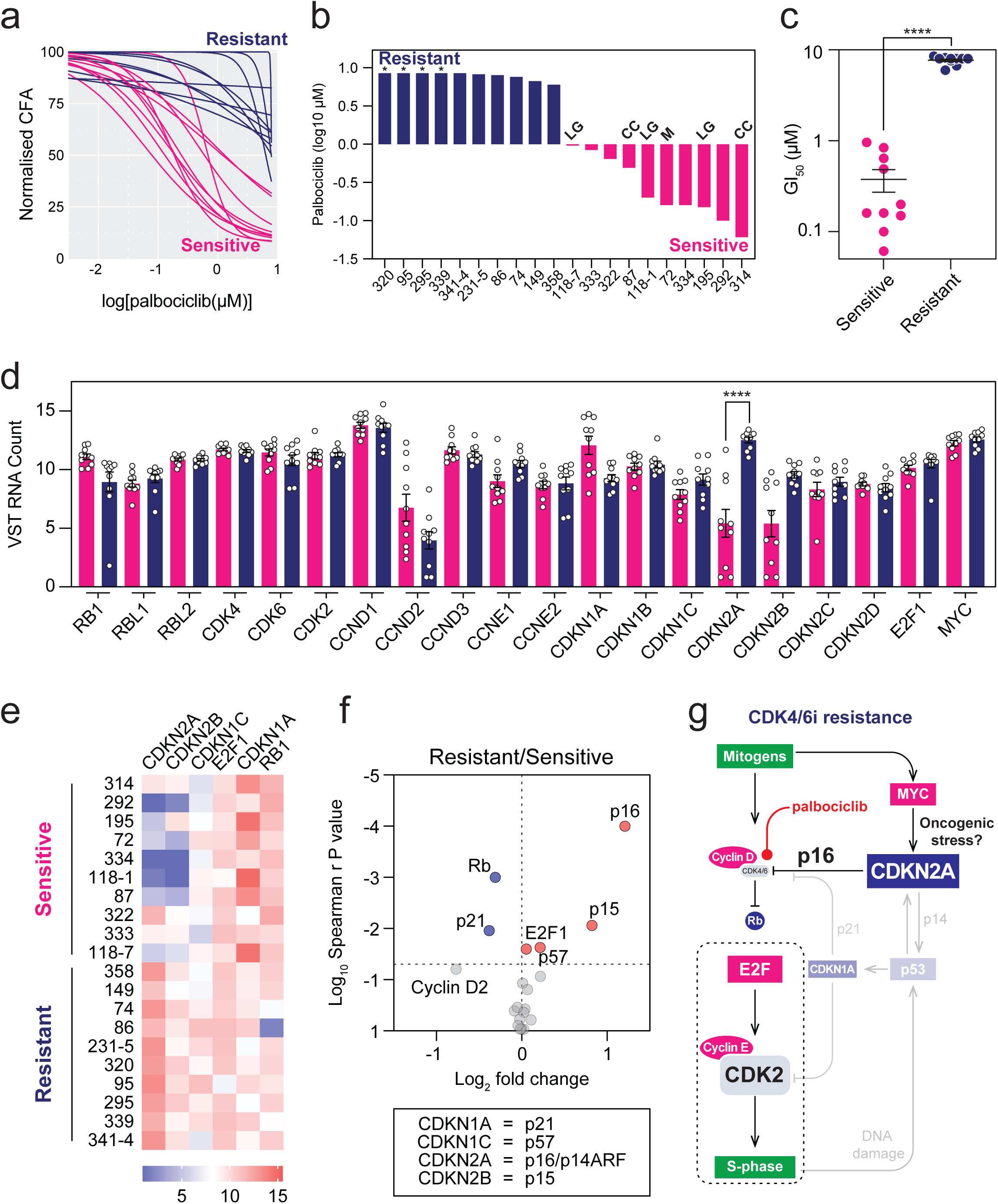
*CDKN2A* is overexpressed in palbociclib-resistant ovarian cancer models a. XY graph showing the dose response curves for all 20 OCMs studied, partitioned into 10 resistant models (dark blue) and 10 sensitive models (magenta). b. Bar graph showing the palbociclib GI_50_ values for 20 OCMs, rank ordered left to right and partitioned into resistant and sensitive models. The six non-HGSOC models are denoted as follows: LG, low grade serous ovarian cancer models; CC, clear cell; M, mucinous. For four resistant HGSOC models (*), accurate GI_50_ values could not be calculated so were assigned nominal values of 8.0 µM. c. Scatter dot pot showing the GI_50_ values for the 20 OCMs. **** p < 0.0001, Mann-Whitney test. d. Bar graph showing normalised transcript reads for 20 G1/S regulator genes indicated, partitioned into resistant (dark blue) and sensitive (magenta) models. Each point represents the RNA-seq derived value for one OCM, such that the bar shows the mean ± s.e.m. derived from 10 data points. For *CDKN2A*, **** p < 0.0001, Kruskal-Wallis test with Dunn’s multiple comparisons test. All other gene comparisons are not significantly different. e. Heatmap showing the relative expression levels of six G1/S regulator genes across 20 OCMs. These six were selected because they have significant Spearman r values when correlating relative mRNA count with GI_50_ values. f. Volcano graph plotting Spearman r P values against fold-change for 20 G1/S regulator genes across 20 OCMs. Key shows the CKI proteins encoded by the indicated genes. Fold-change is calculated as resistant/sensitive, and a p-value threshold of 0.05, shows that p16, p15, p57 and E2F1 are upregulated in resistant OCMs, while Rp and p21 are downregulated. g. Network diagram summarising changes to G1/S cell cycle controls in palbociclib-resistant HGSOC OCMs, whereby upregulation of *CDKN2A* results in p16-mediated inhibition of CDK4/6. As a result of *TP53* mutation, p53-mediated activation of *CDKN1A* is absent, resulting in diminished potency of p21-dependent CDK inhibition. While p16-dependent cellular inhibiton CDK4/6 renders palbociclib ineffective, S-phase entry proceeds due to oncogenic activation of CDK2, via reduced Rb function and/or activation of E2F/Cyclin E.

To further define transcriptional correlates of palbociclib response, we analysed expression data for the 20 G1/S cell cycle control genes across the full 20-OCM panel. In a multiple-comparisons test, only *CDKN2A* was significantly different, with resistant models showing higher *CDKN2A* levels compared with sensitive ones (Figure 4d). Nonetheless, we did observe some additional trends: consistent with our interrogation of the heatmap in Figure 3b, *CDKN2B* tended to be elevated and *RB1* reduced in resistant lines (Figure 4e). In addition, *CDKN1A* and *CCND2*—which encode p21 and cyclin D2, respectively—were lower in resistant models.

To explore this further, we calculated correlation values between palbociclib sensitivity and gene expression, generating a Spearman r P value for each gene. We then averaged the normalised RNA read counts for the sensitive and resistant models, to generate a fold-change value of resistant over sensitive. Plotting – log₁₀(P) against fold-change in a volcano-style graph (Figure 4f) confirmed that *CDKN2A* and *CDKN2B* were significantly upregulated in resistant OCMs (P < 0.05). *E2F1* and *CDKN1C* also reached significance, though with modest fold-changes. Conversely, *CDKN1A* and *RB1* were significantly downregulated in resistant models. Although *CCND2* exhibited a large negative fold-change, it did not achieve statistical significance.

Collectively, these analyses reinforce that low *CDKN2A*/*CDKN2B* and high *RB1* expression correlate with palbociclib sensitivity in OCMs, and suggest this transcriptional signature may predict CDK4/6 inhibitor responsiveness in ovarian cancers representing multiple disease subtypes. Furthermore, it suggests a mechanism whereby palbociclib-resistant OCMs progress through G1 into S-phase despite p16-mediated inhibition of CDK4/6 due oncogenic activation of CDK2 (Figure 4g).

### Diminished palbociclib-sensitivity of a post-treatment low-grade model

Our Living Biobank contains cohorts of longitudinal samples, including subsets with models generated from biopsies collected before, during and after treatment (Nelson et al., 2020; Nelson et al., 2023). For example, for patient 118–who was diagnosed with LGSOC–we collected ascites seven times during the early course of the disease (Figure 5a). Following diagnosis, the patient was treated with carboplatin and paclitaxel, followed by maintenance therapy with the anti-estrogen, letrozole. The post-treatment model, OCM.118-7 featured in our initial screen of 15 OCMs (Figure 2), while the pre-treatment model, OCM.118-1, was selected in the second analysis on the basis of low *CDKN2A/B* expression (Figure 3). To analyse more directly, we compared dose response curves and gene expression profiles side-by-side. Chemo-naïve OCM.118-1 had a palbociclib GI_50_ value of 200 nM, compared with ∼1 µM for OCM.118-7 (Figure 5b), suggesting increasing resistance to CDK4/6 inhibition. This shift was accompanied by a decrease expression of *CDKN2A/B* (Figure 5c). Although each model has only a single RNA-seq profile—precluding formal statistical tests—the parallel changes in GI₅₀ and *CDKN2A/B* expression suggest remodelling of G1/S cell cycle controls during treatment, possibly in response to anti-estrogen therapy.

**Figure 5.**
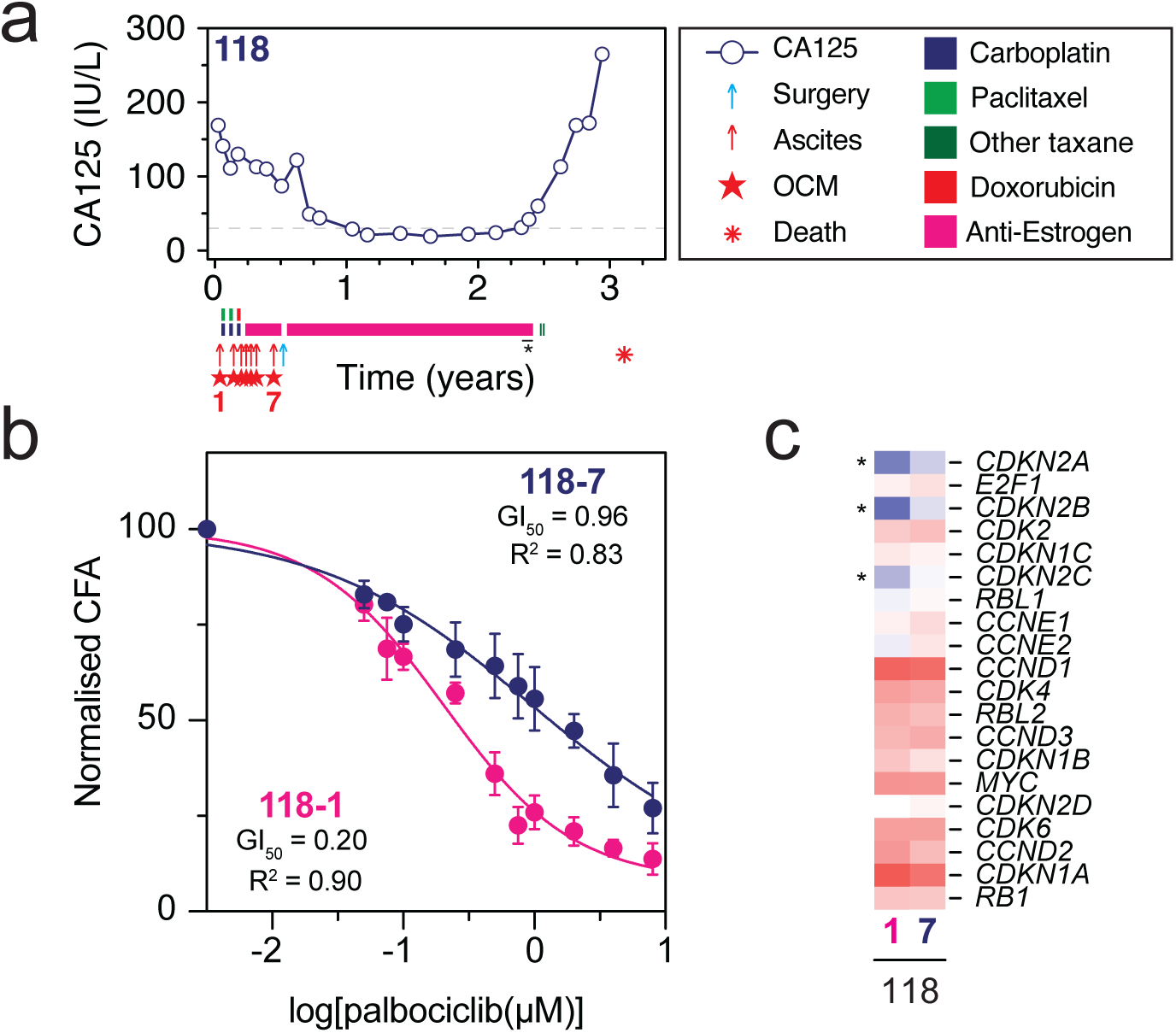
Diminished palbociclib-sensitivity of a post-treatment low-grade model a. Patient Timeline for patient 118 plotting CA125 values (proxy for disease progression) against time in years since being diagnosed with LGSOC, annotated with the treatments received. Blue arrow shows surgery while red arrows show ascitic drains. Red stars indicate samples that gave rise to OCMs, with number below indicating OCMs used in this study. b. XY graph quantitating CFA values across a range of palbociclib concentrations. Each value represents the mean ± s.e.m. from at least three independent experiments. The dose response curve was calculated using a non-linear regression to determine GI_50_ values and goodness of fit (R^2^). c. Heatmap showing the relative expression levels of 20 G1/S regulator for OCMs 118-1 and 118-7, highlighted (*) to show *CDKN2A, CDKN2B and CDKN2C*.

## Discussion

Chemotherapy remains the cornerstone OC treatment; however, responses are rarely durable with HGSOC (Bowtell et al., 2015), and rarer subtypes such as LGSOC, CCOC, and MOC are often intrinsically chemoresistant (Grabowski et al., 2016; Grisham et al., 2023), underscoring the need for alternative therapeutic strategies for these patients. Across these subtypes, deregulation of the G1/S cell cycle controls is prevalent (Singer et al., 2003; Hunter et al., 2015; Friedlander et al., 2016; Cheasley et al., 2019; Cheasley et al., 2021; Iida et al., 2021), including alterations in *RB1*, *CDKN2A*, and *CCNE1*, which create distinct dependencies on specific CDK-cyclin complexes and reveal potential therapeutic vulnerabilities (Knudsen et al., 2022; Zhang et al., 2022; Sine et al., 2025). Yet, these dependencies and vulnerabilities remain poorly defined in OC and preclinical evaluation of CDK inhibitors in clinically relevant OC models are needed. In this study, we screened 20 diverse patient-derived OCMs from our Living Biobank for sensitivity to the CDK4/6 inhibitor palbociclib. We identified a biologically defined subset of models, encompassing HGSOC and rarer subtypes, that was sensitive to palbociclib. These models exhibited low *CDKN2A/2B* expression, and following treatment, inhibition of Rb phosphorylation and induction of G1 arrest. By contrast, resistance was observed in models with consistently high *CDKN2A/*2B expression, low expression of *RB1*–and in one case apparent Rb loss– and *CDKN1A*. Altogether, our findings support a model of G1/S checkpoint control whereby ovarian cancers dependent on CDK4/6–Cyclin D for cell cycle entry exhibit low levels of p16 expression and Rb proficiency (Knudsen et al., 2024). By contrast, in the context of a non-functional p53-p21 axis, those cases not dependent on CDK4/6 are likely driven through downstream activity of CDK2–Cyclin E activity. Importantly, these divergent CDK dependencies give rise to therapeutically actionable vulnerabilities.

Our data indicates that CDK4/6 inhibition may be particularly effective in patients with rarer subtypes such as LGSOC, MOC, and CCOC, diseases that lack effective treatment options. These subtypes were well represented among the palbociclib-sensitive models, consistent with findings in OC cell lines (Konecny et al., 2011). Intriguingly, all chemo-naïve OCMs in our panel were sensitive to palbociclib, raising the possibility that CDK4/6 dependency is an early feature of OC tumour biology that diminishes over time or through selective pressure from treatment. This is exemplified by our longitudinal models from a patient with LGSOC, where chemo-naïve OCM.118-1 exhibited exquisite sensitivity, while the matched post-treatment OCM.118-7 was approximately five-fold more resistant. These findings underscore how CDK-dependencies in OC are dynamic, and highlight the potential value of early intervention with CDK4/6 inhibitors – as illustrated by a case report of a patient with LGSOC who had a marked clinical benefit from frontline anti-oestrogen therapy combined with palbociclib (Handagala et al., 2025). However, two reports from heavily pretreated patients with HGSOC, one with confirmed *CDKN2A* loss, also demonstrated prolonged progression-free and overall survival with palbociclib combined with anti-oestrogen therapy (Frisone et al., 2020; Lee and Ho, 2020). Thus, while CDK4/6 inhibitors may be effective when administered early, selected patients may continue to benefit in later treatment settings, provided favourable molecular features are retained. Indeed, a phase II trial of palbociclib monotherapy in an unselected heavily-pretreated cohort of patients with OC showed only modest clinical activity (Konecny et al., 2016). Another phase II trial combining a CDK4/6 inhibitor with anti-oestrogen therapy demonstrated durable responses in LGSOC, but limited efficacy in HGSOC (Colon-Otero et al., 2020). These clinical findings align with our observations that only a subset of OCMs are sensitive to palbociclib, highlighting the need for biomarker-guided patient selection.

Concordant with previous studies in OC cell lines (Konecny et al., 2011), our study found low *CDKN2A* expression was the strongest individual predictor of palbociclib sensitivity across the OCMs: all resistant OCMs exhibited consistently high *CDKN2A* expression. However, not all sensitive OCMs exhibited low *CDKN2A* expression, suggesting either that *CDKN2A* loss is not essential for CDK4/6-dependency, or that p16 mRNA expression does not fully capture functional protein status. Indeed, p16 is subject to complex regulation (Li et al., 2011), and mRNA-based assessments may overlook post-transcriptional and post-translational dynamics. Similarly, while previous studies report high *CCND1/2* expression correlates with sensitivity, and high *CCNE1/2* with resistance (Konecny et al., 2011; Knudsen et al., 2022; Zhang et al., 2022), we did not observe these associations across the OCMs. Notably, OCM.314 – the most palbociclib-sensitive model in the screen – exhibited the highest *CCNE1* expression of the cohort. These discrepancies highlight the limitations of transcriptomics as standalone biomarkers and underscore the need for more integrated approaches when guiding treatment decisions. To improve patient selection, more robust biomarkers will be required, including for example, p16 immunohistochemistry could be employed at diagnosis to exclude patients unlikely to benefit from CDK4/6 inhibition.

We also examined mechanisms of palbociclib resistance. As all resistant models were HGSOC, where *ABCB1* overexpression is a common chemotherapy resistance mechanism (Patch et al., 2015; Christie et al., 2019), we examined whether drug efflux contributed to palbociclib resistance in the OCMs. Contrary to a report that palbociclib is a *ABCB1* substrate, and *ABCB1* overexpression mediates palbociclib resistance in certain cancer cell lines (Fu et al., 2022), we found that *ABCB1* overexpression did not mediate palbociclib resistance in the OCMs. Pharmacological inhibition of MDR1 with elacridar in the OCMs, or inducible overexpression of *ABCB1* in a palbociclib-sensitive RKO cell line did not affect palbociclib sensitivity, thereby providing no evidence to support MDR1-mediated drug efflux as a major palbociclib resistance mechanism in OC. Instead, resistance in the OCMs is due to cell cycle entry being driven by Cyclin-E-CDK2 activity (Knudsen et al., 2024). This remains speculative: functional validation will be required to confirm whether CDK2 activity mediates palbociclib resistance in the OCMs, for instance through testing the effect of selective CDK2 inhibitors. Indeed, these findings provide a rationale for exploring CDK2 inhibitors in OC. CDK2 inhibitors may offer an alternative strategy for tumours that are intrinsically resistant or acquire resistance to CDK4/6 inhibitors by switching to CDK2 dependency. INX-315, a selective CDK2 inhibitor currently in clinical development having demonstrated remarkable efficacy in CDK2-dependent cell models (Dietrich et al., 2024; Watts and Spencer, 2024; Kumarasamy et al., 2025), has been fast-tracked for *CCNE1*-amplified, platinum-resistant ovarian cancer. Such agents warrant further investigation in palbociclib-resistant OCMs.

In conclusion, this study identifies a clinically and biologically relevant subset of OC, encompassing HGSOC and rarer subtypes, that are dependent on CDK4/6 as evidenced by sensitivity to a selective CDK4/6 inhibitor, particularly those characterised low p16 expression. Our findings support a refined model of G1/S checkpoint control, in which tumours diverge in their reliance on CDK4/6-Cyclin D versus CDK2–Cyclin E for cell cycle entry. This framework provides a molecular rationale for patient selection and highlights the potential of integrating molecular, protein, and functional biomarkers to inform treatment decisions. Longitudinal models such as OCM.118 offer insight into the evolution of CDK dependencies and could guide future therapeutic sequencing. As new selective agents targeting CDK2 become available, mapping both CDK4/6 and CDK2 dependencies and vulnerabilities will be critical to advancing personalised therapies for patients with OC with limited effective, durable treatment options.

### Limitations of the Study

Throughout, assays were performed in 2D cultures of purified tumour cells, which cannot capture the complexity of the tumour microenvironment or stromal cell interactions that may influence cell cycle regulation and drug responses. While colony formation assays captured long-term proliferation responses, their interpretation can be confounded by cytostatic effects (Foy et al., 2024); future studies using GFP-H2B– labelled OCMs with time-lapse imaging could provide more accurate, dynamic measures of proliferation and single-cell behaviour. RNA-seq provided valuable insights but does not account for protein expression or functional interactions. Additionally, the relatively small sample size—particularly for rarer subtypes—and the exclusive use of palbociclib limit the generalisability of the findings, which should be validated across larger cohorts and with other CDK4/6 inhibitors.

## Materials and Methods

### Ovarian Cancer models

Ascitic fluid samples, obtained with informed patient consent from the Manchester Cancer Research Centre (MCRC) Biobank (Human Tissue Authority license: 30004), were used to generate OCMs as previously published (Table S1) (Pillay et al., 2019; Nelson et al., 2020; Barnes et al., 2021; Coulson-Gilmer et al., 2021; Golder et al., 2022; Coulson-Gilmer et al., 2024; Littler et al., 2025; Tighe et al., 2025). The role of the MCRC biobank, which is ethically approved as a research tissue bank by the South Manchester Research Ethics Committee (Ref: 22/NW/0237), is to distribute samples; it does not endorse studies performed or the interpretation of results (see https://www.mcrc.manchester.ac.uk/research/mcrc-biobank for more information). RNA sequencing data associated with the OCMs are published previously (Nelson et al., 2020; Barnes et al., 2021; Coulson-Gilmer et al., 2021; Littler et al., 2025; Tighe et al., 2025).

### Cell culture

Human colorectal cancer cell lines, RKO/LacZeo/TO cells and RKO/FRT/TO/Myc-MDR1 cells expressing GFP-H2B and a tetracycline-inducible Myc-MDR1 construct (Tighe et al., 2025), were cultured in Dulbecco’s Modified Eagle Medium (DMEM) supplemented with 10% foetal bovine serum (FBS), 100 U/ml penicillin, 100 μg/mL streptomycin and 2 mM glutamine and maintained at 37⁰C in a humidified 5% CO_2_ atmosphere. RKO/FRT/TO/Myc-MDR1 cells were selected with 10 mg/ml blasticidin, 50 mg/ml hygromycin B (Sigma Aldrich), and 10 µg/ml G418. OCMs were cultured in Cell+ flasks, except for OCM.86, which was cultured in collagen-coated flasks; containing Ovarian Carcinoma Modified Ince (OCMI) media (Ince et al., 2015) and maintained at 37⁰C in a humidified 5% CO_2_ and 5% O_2_ atmosphere. For long-term storage, RKO cells were cryopreserved in 20% FBS, 10% DMSO, and 70% DMEM, and OCMs cryopreserved in Bambanker (Wako Pure Chemical); and stored at −80⁰C. Cells were periodically tested for the presence of mycoplasma and authenticated (Promega Powerplex 21 System) by the Molecular Biology Core Facility at the CRUK Manchester Institute.

### Drugs

Palbociclib (Santa Cuz Biotechnology), elacridar (Selleckchem), and paclitaxel (Sigma-Aldrich) were dissolved in DMSO. Tetracycline was dissolved in H_2_O. All agents were aliquoted stored at −80⁰C or −20⁰C.

### Colony Formation Assay

Cells were seeded into 12-well plates at 250 cells/well for RKO cell lines, or 500–5000 cells/well for OCMs; and incubated overnight. A titration of palbociclib (0, 50, 75, 100, 250, 500, 750, 1000, 2000, 4000, 8,000 nM) and 3.2% (v/v) DMSO was added to the cells. Cells were incubated with drugs for up-to 21 days, with media containing drugs replaced weekly. For some OCMs, additional CFAs using a lower titration of palbociclib were performed (0, 5, 10, 25, 75, 100, 250, 500, 750, 1000 nM). CFAs were performed with a palbociclib titration in combination with 250 nM elacridar as indicated. For RKO/FRT/TO/Myc-MDR1 CFA, cells were plated in the presence or absence of 1 μg/mL tetracycline, and the following day, a titration of palbociclib with or without 1 μg/mL tetracycline was added.

Once a single well had reached confluence, or 21 days post-treatment, cells were fixed in 1% (v/v) formaldehyde (Fisher Scientific) for 10 min, stained with a 0.05% (v/v) crystal violet (Sigma-Aldrich) for 10 min, and imaged on a ChemuDoc Touch Imaging System (BioRad). Crystal violet was extracted with 10% (v/v) acetic acid and absorbance at 570 nm read on a VarioSkanLUX multimode microplate plate reader (Thermo Scientific). The mean absorbance from six 10% (v/v) acetic acid-only blank wells was subtracted from each value, before normalisation to the DMSO control well. Data from at least three biological replicates were used to generate dose-response curves and determine GI_50_ values (concentration at 50% maximal inhibition of growth) using Prism 10 (GraphPad).

### Immunoblotting

Following 24 h treatment with 1 μM palbocilcib or 0.4% (v/v) DMSO, cells were harvested by scrapping, centrifuged at 1,000 rpm for 5 min at room temperature (RT), and pellets stored at −20⁰C. Cells were lysed with RIPA buffer and protein concentration quantified by Bradford assay. Lysates were boiled in SDS sample buffer (0.35 M Tris pH 6.8, 0.1 g/ml sodium dodecyl sulphate, 93 mg/ml dithiothreitol, 30% (v/v) glycerol, 50 μg/ml bromophenol blue; all from Sigma Aldrich) and 20 μg protein resolved by SDS-PAGE using NuPAGE^TM^ 4– 12% (v/v) Bis-Tris protein gels (Life Technologies), before electroblotting onto methanol-soaked Immobilon-P PVDF membranes (Merck Millipore). Membranes were blocked in either 5% (w/v) dried skimmed milk (Marvel) or 5 % (w/v) bovine serum albumin (BSA, Sigma Aldrich) (for phospho-antibody) dissolved in TBS-T (50 nM Tris pH 7.6, 150 mM NaCl, 0.1% Tween-20), and incubated at 4⁰C overnight with primary antibodies: mouse anti-Rb (9307L Cell Signalling Technology, 1:2,000), rabbit anti-phospho-Rb serine-780 (9309T, Cell signalling Technology; 1:2,000) and mouse anti-TAT1 (1:2,500; a kind gift from I Hagan). Membranes were washed in TBS-T and incubated with horseradish-peroxidase (HRP)-conjugated secondary antibodies: goat anti-mouse IgG [H+L] HRP, Invitrogen, cat#G21040 RRID: AB_2536527; goat anti-rabbit IgG [H+L] HRP, Invitrogen, cat#G21234, RRID: AB_1500696; for 2 h. After TBS-T washes, bound secondary antibodies were detected using EZ-Chemiluminescence Reagent (Sartorius) and a ChemiDoc^TM^ Touch Imaging System (BioRad). At least two biological replicates were performed.

### Flow Cytometry

For DNA content analysis, cells were exposed to 1 μM palbociclib or 0.4% (v/v) DMSO for 24 h, conditioned media collected, cells harvested by trypsinisation, samples pooled and centrifuged at 1,000 rpm for 5 min at room temperature. Cell pellets were fixed in 70% (v/v) ice-cold ethanol and stored at −20⁰C for >16 h. Fixed cells were washed in cold PBS and centrifuged at 1500 rpm at 4⁰C for 5 min, twice, and stained with propidium iodide (PI) solution (0.4% (w/v) PI, 0.5% (w/v) RNase A in PBS) for >30 min in the dark at room temperature. Cells were filtered, analysed on an Attune^TM^ NxT flow cytometer (ThermoFisher) with 30,000 events collected per sample using Attune NxT Software. Singlets were gated and cell cycle profiles analysed using FlowJo software (FlowJo, LLC). Two independent biological replicates were performed.

### Time-lapse Microscopy Drug Profiling Assay

RKO/FRT/TO/Myc-MDR1 cells expressing GFP-H2B and a tetracycline-inducible Myc-MDR1 construct were seeded into black μclear® 96-well plates (Greiner Bio-One) at 5×10^4^ cells/mL in the presence or absence of 1 μg/ml tetracycline, prior to drug addition. Plates were maintained at 37⁰C in a humidified 5% CO_2_ atmosphere and imaged using an IncuCyte® S3 (Sartorius AG) with a 20x objective. Images were acquired every 4 h for 168 h. IncuCyte® S3 software was used in real time to measure green object count as a proxy for proliferation. Green object counts were normalised to time t=0 and proliferation curves generated. Two biological replicates were performed.

### Statistical Analysis

Microsoft Excel was used for data organisation and Prism 10 (GraphPad) for statistical analysis, where ∗*p* < 0.05, ∗∗*p* < 0.01, ∗∗∗*p* < 0.001, ∗∗∗∗*p* < 0.0001, ns: *p* > 0.05. Details of statistical analyses are described in the figure legends.

## DATA AVAILABILITY

The gene expression data presented here has been deposited with EMBL-EBI, accession numbers: E-MTAB-7223; E-MTAB-10801; E-MTAB-11000; E-MTAB-14568

## AUTHOR CONTRIBUTION STATEMENT

McDonald-Pike: conceptualisation, formal analysis, methodology, visualisation, and writing–original draft. Coulson-Gilmer, Littler, Barnes, Altringham and Tighe: methodology, resources, and supervision. McGrail: writing–review and editing and project administration. Taylor: conceptualisation, funding-acquisition, validation, visualization, writing–review & editing.

## ACKNOWLEDGEMENTS

Research in the Taylor lab is funded by Cancer Research UK [C1422/A31334] and [C147/A25254]; the Medical Research Council [MR/X008088/1]; and the NIHR Manchester Biomedical Research Centre [NIHR203308]; the views expressed are those of the author(s) and not necessarily those of the NIHR or the Department of Health and Social Care. We thank members of the Taylor lab for experimental advice and comments on the manuscript.

## Supplementary Information

**Table S1.** Patient and OCM characteristics.

| Patient |  |  |  | OCM |  |  |  |  |  |  |
| --- | --- | --- | --- | --- | --- | --- | --- | --- | --- | --- |
| # | ID | FIGO Stage | Subtype<sup>a</sup> | OCM ID<sup>b</sup> | Chemo-naïve (Y/N)<sup>c</sup> | TP53<br>(sequencing of cloned transcripts) |  | p53 immunostaining<br>+/- Nutlin-3 |  | References |
|  |  |  |  |  |  | DNA | Protein prediction | -Nutlin-3<sup>d</sup> | +Nutlin-3<sup>e</sup> |  |
| 1 | 72 | 1A | Moderately differentiated MOC | 72 | N | c.843C>G | p.D281E | + | - | 2, 3, 5, 7, 8 |
| 2 | 74 | 3B | HGSOC | 74 | N | c.1024C>T | p.R342X | + | - | 2, 3, 4, 8 |
| 3 | 86 | 4B | HGSOC | 86 | N | c.842A>G | p.D281G | - | - | 3, 4, 7, 8 |
| 4 | 87 | 3B | Possible CCOC<sup>†</sup> | 87 | Y | WT | WT | + | + | 2, 3, 4, 5, 7, 8 |
| 5 | 95 | 3C | HGSOC | 95 | N | c.902del | Frameshift | + | - | 4, 8 |
| 6 | 118 | 3C | LGSOC | 118-1 | Y | ND | ND | - | + | 1, 3, 8 |
|  |  |  |  | 118-7 | N | ND | ND | - | + | 3, 8 |
| 7 | 149 | 3C | HGSOC | 149 | N | c.724T>G | p.C242G | + | - | 3, 4, 7, 8 |
| 8 | 195 | 4A | Possible LGSOC<sup>†</sup> | 195 | Y | WT | WT | + | + | 3, 4, 5, 7, 8 |
| 9 | 231 | 3C | HGSOC | 231-5 | N | c.742C>T | p.R248W | + | - | 7, 8 |
| 10 | 292 | 3C | HGSOC | 292 | N | c.63delC | fs. Stop 43 | - | - | 8 |
| 11 | 295 | 3C | HGSOC | 295 | N | c.757_758insA | Frameshift | - | - | 8 |
| 12 | 314 | 1A | CCOC | 314 | Y | WT | WT | - | + | 8 |
| 13 | 320 | 3C | HGSOC | 320 | N | c.742C>T | p.R248W | + | - | 8 |
| 14 | 322 | 3A | HGSOC | 322 | N | c.400T>G | p.F134V | + | - | 8 |
| 15 | 333 | 3C | HGSOC | 333 | N | c.524G>A | p.R175H | - | - | 8 |
| 16 | 334 | 3C | HGSOC | 334 | Y | c.743G>T | p.R248L | + | - | 8 |
| 17 | 339 | 4A | HGSOC | 339 | N | c.527G>A | p.C176Y | + | - | 8 |
| 18 | 341 | 3C | HGSOC | 341-4 | N | c.524G>A | p.R175H | - | - | 9 |
| 19 | 358 | 3C | HGSOC | 358 | N | ND | ND | - | - | 8 |

**Figure S1.**
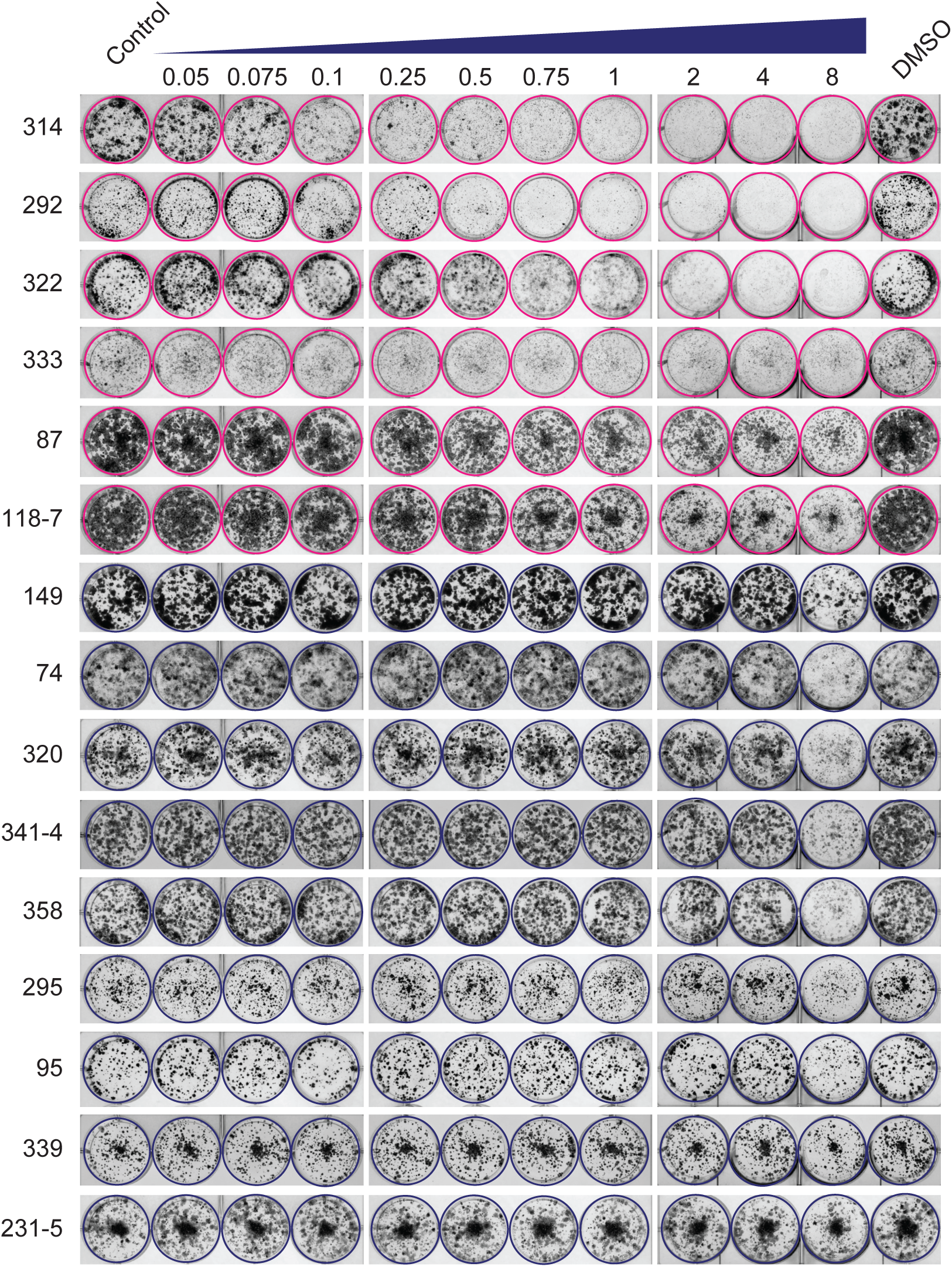
Palbociclib inhibits a subset of patient-derived ovarian cancer models Representative colony formation assays for the 15 ovarian cancer models screened for palbociclib sensitivity which identified six sensitive models (magenta) and nine resistant models (dark blue).

**Figure S2.**
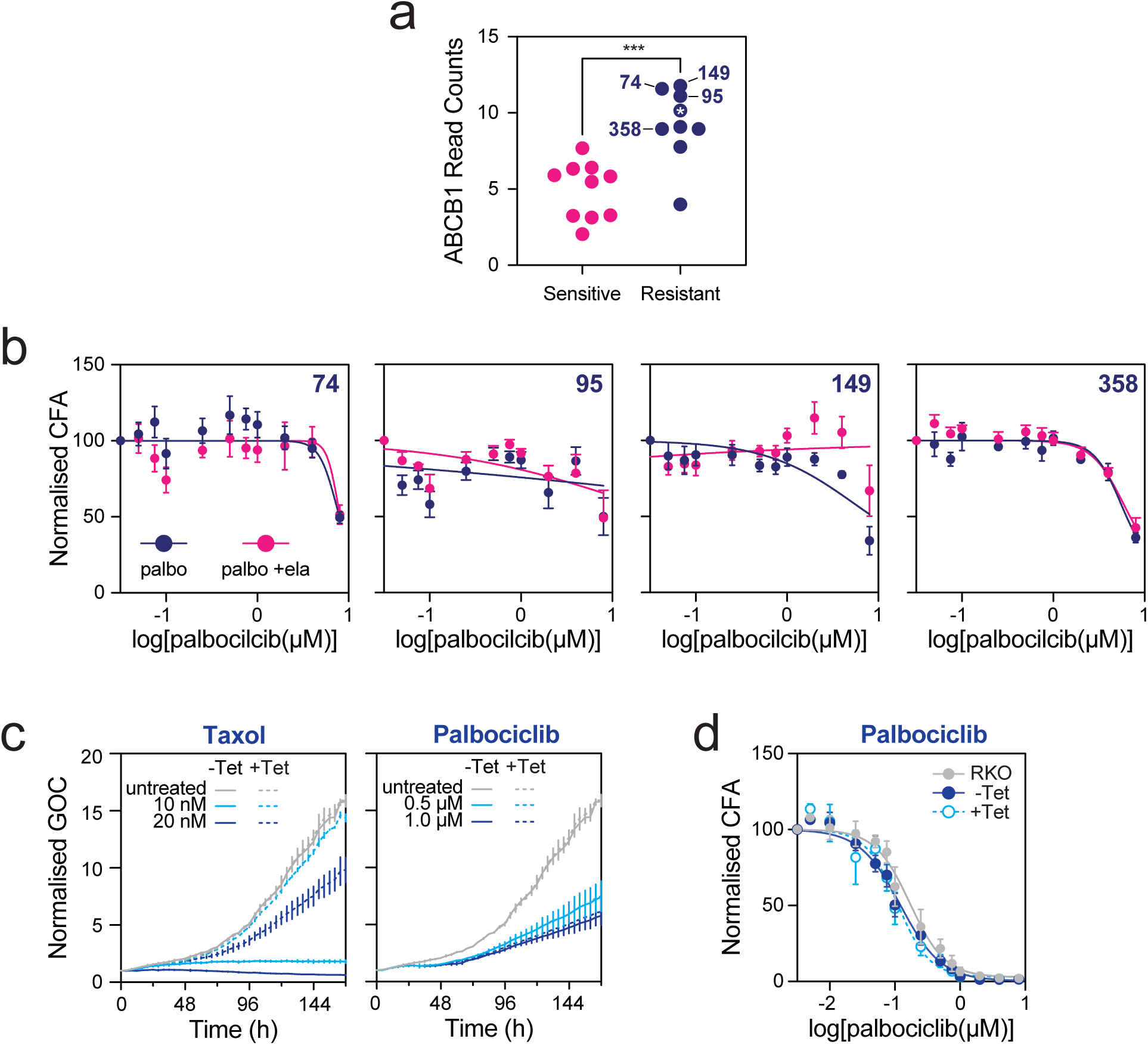
Inhibition of MDR1-depedent drug eflux does not sensitise palbociclib-resistant ovarian cancer models a. Scatter dot pot showing the normalised *ABCB1* read counts for the 20 OCMs partitioned into resistant (dark blue) and sensitive models (magenta). *** p < 0.001, Mann-Whitney test. White asterisk (*) highlights OCM.320 which overexpresses *ABCB1* transcript but does not overexpress MDR1 protein (A.Tighe, unpublished). b. XY graph quantitating CFA values for four OCMs with relatively high *ABCB1* expression, across a range of palbociclib (palbo) concentrations, with and without the MDR1 inhibitor, elacridar (ela). Each value represents the mean ± s.e.m. from at least three independent experiments. c. Proliferation curves plotting normalised green object count (GOC) against time in hours for RKO cells expressing a GFP-tagged histone. Cells were exposed to either taxol or palbociclib at the concentrations indicated, plus/minus tetracycline to induce expression of an MDR1 transgene. d. XY graph quantitating CFA values for RKO cells in response to a range of palbociclib concentrations. RKO cells are either the parental line (grey), or a derivative harbouring a tetracycline inducible MDR1 transgene, analysed in the absence (dark blue) or presence (cyan) of tetracycline.

**Figure S3.**
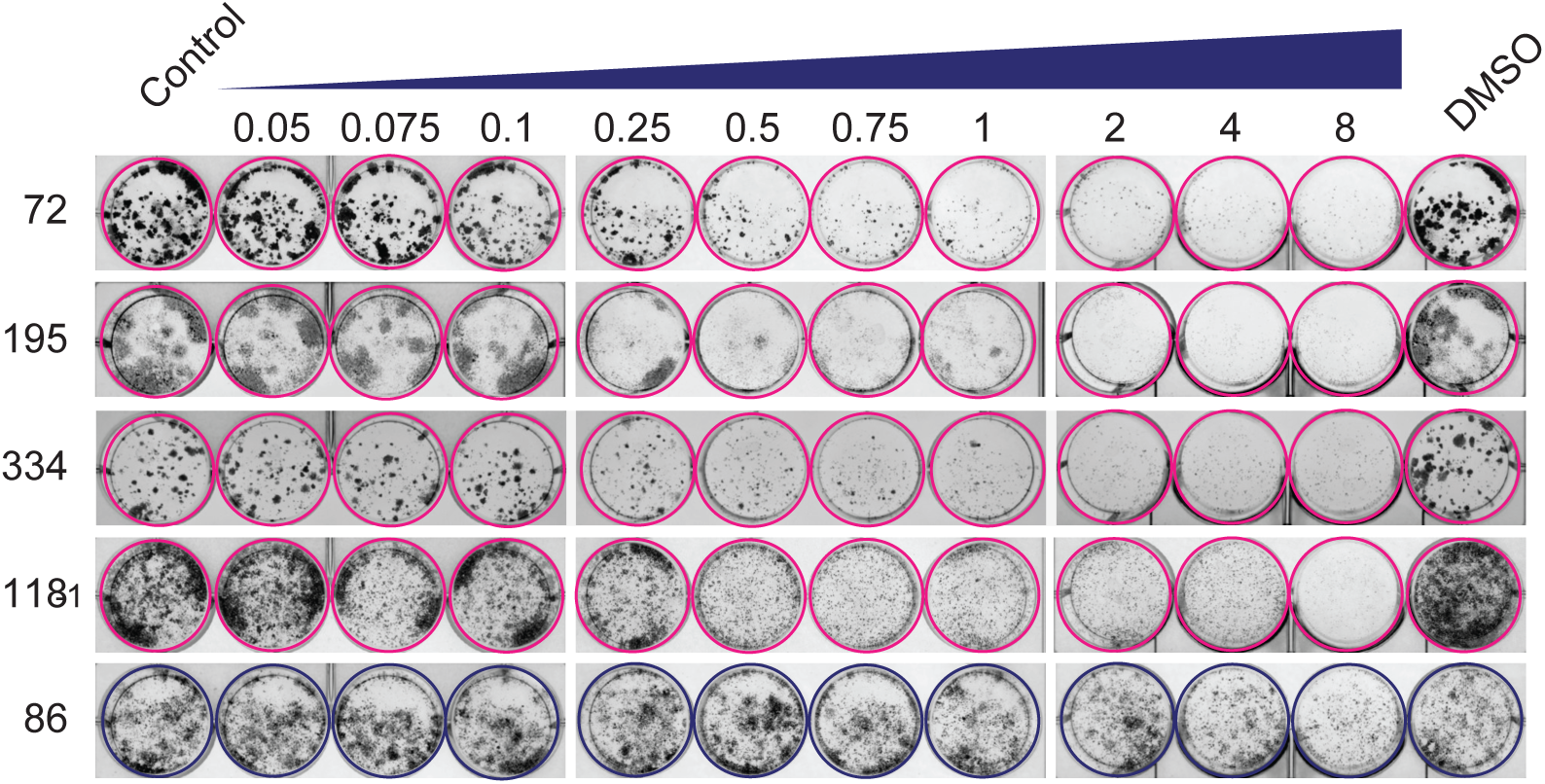
Identification of additional palbociclib-resistant models Representative colony formation assays for the five ovarian cancer models screened for palbociclib sensitivity, identifying a further four sensitive models (magenta) and one resistant model (dark blue).

